# Genome improvement and genetic map construction for *Aethionema arabicum*, the first divergent branch in the Brassicaceae family

**DOI:** 10.1101/662684

**Authors:** Thu-Phuong Nguyen, Cornelia Mühlich, Setareh Mohammadin, Erik van den Bergh, Adrian E. Platts, Fabian B. Haas, Stefan A. Rensing, M. Eric Schranz

**Author notes:** EMBL-EBI, Wellcome Genome Campus, Hinxton, CB10 1SD, United Kingdom.

## Abstract

**Background:** The genus *Aethionema* is a sister-group to the core-group of the Brassicaceae family that includes *Arabidopsis thaliana* and the Brassica crops. Thus, *Aethionema* is phylogenetically well-placed for the investigation and understanding of genome and trait evolution across the family. We aimed to improve the quality of the reference genome draft version of the annual species *Aethionema arabicum*. Secondly, we constructed the first *Ae. arabicum* genetic map. The improved reference genome and genetic map enabled the development of each other.

**Results:** We started with the initially published genome (version 2.5). PacBio and MinION sequencing together with genetic map v2.5 were incorporated to produce the new reference genome v3.0. The improved genome contains 203 MB of sequence, with approximately 94% of the assembly made up of called bases, assembled into 2,883 scaffolds. The N_50_ (10.3 MB) represents an 80-fold over the initial genome release. We generated a Recombinant Inbred Line (RIL) population that was derived from two ecotypes: Cyprus and Turkey (the reference genotype. Using a Genotyping by Sequencing (GBS) approach, we generated a high-density genetic map with 749 (v2.5) and then 632 SNPs (v3.0) was generated. The genetic map and reference genome were integrated, thus greatly improving the scaffolding of the reference genome into 11 linkage groups.

**Conclusions:** We show that long-read sequencing data and genetics are complementary, resulting in an improved genome assembly in *Ae. arabicum*. They will facilitate comparative genetic mapping work for the Brassicaceae family and are also valuable resources to investigate wide range of life history traits in *Aethionema*.

## Background

The genus *Aethionema* belongs to the important plant family Brassicaceae. The crucifers contain many species of interest, such as the Brassica crop plants (e.g. *B. rapa, B. oleracea* and *B. napus*), ornamental plants (such as the genera *Aubrieta, Iberis, Lunaria* and *Draba*) and research model plant species (including *Arabidopsis thaliana, A. lyrata, Capsella rubella* and *Arabis alpina*). Phylogenetic studies have established *Aethionema* as the sister-group of the core-group in the family [1]. Thus, *Aethionema* holds an essential phylogenetic position for studies on genome and trait evolution across the Brassicaceae family.

The monogeneric tribe *Aethionemeae* consists of 57 species and is distributed mainly in the Irano-Turanian region, a hot spot for species radiation and speciation [2–4]. This tribe displays various interesting morphological and ecological characteristics, especially fruit and seed heteromorphism. Heteromorphism is defined as the production of two or more distinct fruit or seed morphs on the same individual [5], which includes morphological size, shape and color; physiological dormancy and germination of fruits and seeds. *Aethionema arabicum* is one of the seven reported heteromorphic species of *Aethionema* [6, 7]. *Aethionema arabicum* is a small diploid annual, with a short life cycle starting from seed germination to the end of the vegetative development in spring, followed by reproduction and the end of life cycle in summer [8]. Both annual life history and heteromorphism probably evolved as adaptive responses to unpredictable environments, especially dry arid habitats, indicating a wide range of natural variation for ecologically adaptive traits in *Ae. arabicum*.

Owing to its unique phylogenetic position and interesting characteristics, *Ae. arabicum* is an ideal sister-group model for research. Therefore, *Aethionema* genome and genetic resources are desirable. The initially published *Ae. arabicum* draft genome (v1.0) contains 59,101 scaffolds with an N50 of 115,195 bp while the genome was predicted to be 320 Mbp in size with n=11 [9]. Here we first aimed to (i) improve the quality of the reference genome and (ii) to construct the first *Ae. arabicum* genetic map. A higher quality version of the genome assembled by the VEGI consortium was later released as version 2.5, which is used as the starting point of our analyses.

High throughput sequencing using Pacific Biosciences (PacBio) and Oxford Nanopore MinION (MinION) technology followed and resolved many uncalled gaps in the v2.5 genome and supported further super-scaffolding, which resulted in genome v3.0. The genetic map was constructed using Genotyping by Sequencing (GBS) on a Recombinant Inbred Line (RIL) population. The 216 RILs were derived from a cross between Turkey (reference ecotype) and Cyprus ecotypes. The first version of genetic map v2.5 was obtained based on genome v2.5 with 746 Single Nucleotide Polymorphism (SNP) markers. The later genetic map v3.0 was built with 626 SNPs generated based on genome v3.0.

Here we show that the long-read genome assembly and the genetic map of *Ae. arabicum* supported the development of each other. They will serve as a substantial resource for further research on *Aethionema* as well as the Brassicaceae family.

## Data description and Methods

### Overview of the workflow

An overview of the improvements of the genome of *Aethionema arabicum* and the generation and improvement of its genetic map are depicted in Figure1. The genome draft version 1.0 was first improved by Ray [10] and AllPathsLGs [11] and led to the release of genome v2.5 (available on genomevolution.org). Genome v2.5 was used as a basis for SNP calling after GBS of the RILs. This generated SNP markers used to construct the genetic map v2.5. Scaffolds were ordered with AllMaps [12] based on the maximum co-linearity to genome v2.5 and genetic map v2.5. This resulted in genome vAM. Gap filling and super-scaffolding improvement for genome vAM was obtained by PacBio sequencing leading to genome v2.6. PBjelly2 [13] run using the MinION reads further improved genome v2.6 to v3.0. We revisited the genetic map v2.5 by recalling SNPs according to genome v3.0 and constructed a genetic map v3.0 with the newly called SNP markers. Below we describe the workflow in detail in the three following sections: (i) the initial genome assembly, (ii) genetic map construction and (iii) genome improvement.

### (i) The intial genome (v2.5): The starting point

#### Genome re-assembly using AllPathsLG

The version 1.0 assembly generated by the Ray assembler [10] was fragmented *in silico* into a set of artificial overlapping reads, combined with paired end and mate pair data (described in [9]), and re-assembled using the AllPathsLG assembler [11]. Gap closing was then performed using GapCloser, part of the SOAPdenovo2 package [14]. Gene annotations were lifted over from assembly version 1.0 to version 2.5 using the LiftOver tool from the UCSC Genome Browser tools package [15].

Genome version 2.5 contains 3,166 scaffolds, has an N50 of 564,741 bp and was published as version 2.5 on https://genomevolution.org/coge/.

### (ii) Genetic map construction

#### Plant material

Two *Aethionema arabicum* ecotypes were used, Turkey (TUR) and Cyprus (CYP). The TUR accession comes from the living plant collections at the Botanical Garden in Jena, Germany (Botanischer Garten Jena). The seeds for this genotype were derived from a plant in the Botanical Garden in Nancy, France. The CYP ecotype was collected in 2010 near Kato Moni (coordinates UTM WGS 84: 508374 - 3879403) at an altitude 410 m on pillow lava by Charalambos S Christodoulou [16].

These two ecotypes were used as parents for the development of the recombinant inbred line population, where TUR was the father and CYP the mother. Seeds from initial F_1_ plants were used to generate an initial F_2_ population. For each of the 216 segregating F_2_ plants, a single seed was randomly chosen to further grow and reproduce the next generation. The procedure was repeated until F_8_, when the experiment was performed with 216 RILs.

To grow the plants for the GBS, F_8_ seeds of 216 RILs were placed on filter paper, wetted with distilled water, in petri dishes. Imbibed seeds were incubated at 4°C in dark for 3 days, followed by germination in the light at 20°C for 2 days. Seedlings were transferred to soil pots (10.5 cm diameter 10 cm height) in November 2014. Plants were grown in greenhouse (Wageningen University and Research, the Netherlands) in partially controlled conditions, long day (16 h light and 8 h dark) and at 20°C.

#### Genotyping By Sequencing (GBS)

##### DNA isolation

Young tissues from leaves and flower buds were collected from each F9 plant for DNA isolation. The DNA isolation was done according to a modified CTAB protocol [17]. In brief, plant material was frozen with liquid nitrogen and ground into powder. Each sample was incubated with 500 µl of CTAB buffer at 65°C in the water bath for 30 min. After 30 min cooling at room temperature, equal volume (500 µl) of chloroform:isoamylalcohol (24:1 v/v) was added, and vigorously hand-mixed for a min. 400 µl of supernatant was recovered after centrifuging at maximum speed for 5 min. The supernatant was cleaned again with a chloroform:isoamylalcohol step. DNA precipitation was performed by adding an equal volume of cold isopropanol with 30 min incubation on ice and centrifugation at maximum speed for 15 min. The DNA pellet was cleaned twice with 1 ml of 70% ethanol and centrifugation at maximum speed for 5 min. Dry DNA pellet was dissolved in Milli-Q water.

##### Constructing GBS libraries

DNA was treated with RNAse overnight at 37°C with RNAse one by Promega. Quality was checked on a 1% agarose gel and DNA quantity was checked with Pico Green. Based on this, DNA was diluted down to 20 ng/µl with MQ water and used in further analysis. GBS was performed in general by following the procedure described in [18]. Oligonucleotides for creation of common as well as 96 barcoded ApeKI adapters were obtained from Integrated DNA Technologies and diluted to 200 µM. For each barcoded and common adapter, top and bottom strand oligos were combined to a 50 µM annealing molarity in TE to 100 µl total volume. Adapter annealing was carried out in a thermocycler (Applied Biosystems) at 95°C for 2 min, ramp to 25°C by 0.1 degree per second, hold at 25°C for 30 min and 4°C forever. Annealed adapters were further diluted to a 0.6 ng/µl concentrated working stock of combined barcoded and common adapter in 96 well microtiter plate and dried using a vacuum oven. For each genomic DNA sample 100 ng (10 ng/µl) was used and added to lyophilized adapter mix and dried down again using a vacuum oven.

Adapter DNA mixtures were digested using 2.5 Units ApeKI (New England Biolabs) for 2 hours at 75°C in a 20 µl volume. Digested DNA and Adapters were used in subsequent ligation by 1.6 µl (400 Units/µl) T4DNA Ligase in a 50 µl reaction volume at 22°C for one hour followed by heat inactivation at 65°C for 30 min. Sets of 96 digested DNA samples, each with a different barcode adapter, were combined (10µl each) and purified using a Qiaquick PCR Purification columns (Qiagen). Purified pooled DNA samples were eluted in a final volume of 10µl. DNA Fragments were amplified in 50 µl volume reactions containing 2 µl pooled DNA, 25 µl KAPA HiFi HotStart Master Mix (Kapa Biosystems), and 2 µl of both PCR primers (12.5 µM). PCR cycling consisted of 98°C for 30 seconds, followed by 18 cycles of 98°C for 30 seconds, 65°C for 30 seconds, 72°C for 30 seconds with a final extension for 5 minutes and kept at 4°C. Amplified libraries were purified as above but eluted in 30 µl. Of the amplified libraries 1 µl was loaded onto a Bioanalyzer High Sensitivity DNA Chip (Agilent technologies) for evaluation of fragment sizes and 1 µl was used for quantification using Qubit (Life Technologies). Amplified library products were used for extra size selection using 2% agarose gel cassette on a blue pippin system (Sage Science) to remove fragments smaller than 300 bp. Eluted size selected libraries were purified by AmpureXP beads (Agencourt). Final libraries were used for clustering on five lanes of an illumina Paired End flowcell using a cBot. Sequencing used an illumina HiSeq2000 instrument using 2*100 nt Paired End reads.

##### Sequencing and processing raw GBS data

Raw sequencing data was processed using the TASSEL software package [19] version 5.2.37 using the GBSv2 pipeline. For quality filtering and barcode trimming, the GBSSeqToTagDBPlugin was run with the following parameters: kmerLength: 64, minKmerL: 20, mnQs: 20, mxKmerNum 100000000. Tags were dumped from the produced database using TagExportToFastqPlugin and mapped to the reference genome using the bwa software package [20] in single-ended mode (samse). Positional information from aligned SAM files was stored in the TASSEL database using the SAMToGBSdbPlugin. The DiscoverySNPCallerPlugin was run using the following parameters: mnLCov: 0.1, mnMAF: 0.01. Found SNPs were scored for quality using SNPQualityProfilerPlugin and the Average taxon read depth at SNP was used as a quality score for filtering in the next step (minPosQS parameter), these scores were written to the TASSEL database using UpdateSNPPositionQualityPlugin. Finally, the ProductionSNPCallerPluginV2 was run with the following parameters: Avg Seq Error Rate: 0.002, minPosQS: 10, mnQS: 20.

##### Genetic map calculation

We used JoinMap v4.1 for the genetic map construction [21, 22]. The genetic map v2.5 was built with 749 SNPs generated by GBS based on genome v2.5 (unprocessed and processed data available as S1 and S2). A set of 632 SNPs called according to genome 3.0 was used for the genetic map v3.0 (unprocessed and processed data available as S3 and S4). Regression and Maximum likelihood mapping were used to calculate these maps (the linkage group information for both 2.5 and 3.0 genetic maps are available as S5).

### (iii) Genome improvement

#### Genome version vAM: AllMaps

We ran AllMaps [12] with default setting to combine genetic map v2.5 and physical map genome v2.5. This step resulted in genome vAM, in which scaffolds were ordered and oriented to reconstruct chromosomes.

#### Contamination removal

The *Ae. arabicum* scaffolds v2.5 were checked for contaminations. The genome scaffolds were split into 197,702 1 kbp fragments and blasted against the NCBI nt database [23] using Tera-BLAST (TimeLogic® Tera-BLAST™ algorithm, Active Motif Inc., Carlsbad, CA). The output was then analyzed by MEGAN 6 [24]. All scaffolds for which more than 50% of their entire length was found in bacteria and with no hit in Viridiplantae were marked as putatively contaminated. A hit was counted with a minimum bit score of 50. Additionally, Bisulfite sequencing (BS-seq) CpG and Chromatin ImmunoPrecipitation DNA-Sequencing (ChIP-seq) H3 data were checked to identify contaminants (Aethionema_contamination.xlsx). We ensured that these scaffolds were not combined with another scaffold by PBjelly2. After screening, three v2.5 scaffolds were removed: Scaffold_2406, Scaffold_2454 and Scaffold_2594. They had a total length of 1,758 bp without any annotated genes. A summary of the contamination screen is available as S6.

#### Long read generation for genome improvement

##### PacBio reads

Genomic DNA (gDNA) for *Ae. arabicum* was obtained from leaves of the Cyprus and Turkey ecotypes. DNA was extracted using a modified protocol [25] based on [26]. For the Turkey ecotype 35.70 µg and for the Cyprus ecotype 21.45 µg high molecular weight DNA were sent to the Max Planck-Genome-Centre, Cologne, and sequenced using the PacBio RS II machine (library insert size was 10-15 kbp gDNA). Four flow cells for Cyprus and six for Turkey were sequenced. Table 1 summarizes the statistics of the reads. The CG content of the pooled reads was 38%.

**Table 1:** Overview of the *Aethionema arabicum* PacBio reads.

|  | Total reads | Cyprus | Turkey |
| --- | --- | --- | --- |
| Number of reads | 381,069 | 152,415 | 228,654 |
| Length variation | 11 - 57,910 | 11 - 55,919 | 11 - 57,910 |
| Average length | 5,845.12 | 5,794.88 | 5,878.8 |
| Average quality | 10.48 | 10.13 | 10.72 |

The lengths are given in nucleotides and the quality as phred score.

##### MinION reads

To obtain MinION reads, gDNA was extracted from the Turkey ecotype (four leaf samples, 100 mg each) as outlined above. After pooling the samples, the gDNA concentration was measured using Hoechst 33258 DNA dye and resulted in 73.85 ng/µl. The library preparation was done using the Oxford Nanopore SQK-NSK007 protocol and R9.4 chemistry to design a 8 kbp 2D library. The sequencing run was carried out using Oxford Nanopores MinION technology. The flow cell sequenced 30,935 reads (122,362,072 nt) at −205 mV and 24 hours of runtime. After base calling with the MinKNOW 1.6 software (Oxford Nanopore Technologies Ltd.) the read length ranged from 5 to 63,441 nt with an average length of 3,955 nt. The average phred quality score was 11 and the GC content 41%, reads were not filtered or trimmed. The initial sequence format FAST5 was converted to FASTQ format by using the R package poRe version 0.21 [27]. Because the MinION flow cell had previously been used for *Physcomitrella patens* DNA in the same run, the 30,935 reads were filtered for putative *P. patens* contamination. The reads were mapped with the long read mapper GMAP version 2017-08-15 [28] against the *P. patens* genome V3 [29]. All settings were kept at default. 1,447 reads were characterized as putative *P. patens* reads and therefore removed.

#### Genome improvement using long reads

To perform super-scaffolding and gap filling, the program PBjelly 2 version 15.8.24 was used [13]. It internally uses BLASR v5.1 [30] for mapping reads to a reference. The BLASR parameters internally used for mapping were: “-minMatch 12 -bestn 1 - noSplitSubreads”.

#### Genome version 2.6: PacBio sequencing incorporation

We ran PBjelly2 with 381,022 (152,398 CYP, 228,624 TUR) PacBio reads which where head-cropped with 20 (due to suspicious per base sequence content suggestion presence of adapters) using Trimmomatic version 0.36 [31].

PBjelly2 was used to improve genome v2.5 and vAM. Comparing the results, we found some scaffold connections which were made by PBjelly2 (v2.5) were no longer possible for vAM (these scaffolds were already connected). Five connections formed for v2.5 scaffolds were already introduced by the genetic map approach (see above). Twelve connections which could be established in v2.5 were not formed in the PBjelly2 output for improving vAM, because the scaffolds were already connected with other scaffolds. Since PBjelly2 only fills gaps with reads and is not able to place whole scaffolds in gaps, it was necessary to split the vAM genome at certain points to be able to obtain the twelve connections which were not present in the PBjelly2 output for vAM (visualized in Figure 2). Split scaffolds were reconnected again after running PBjelly2, using N-stretches of length 100 to keep all improvements introduced in vAM if they were not formed by PBjelly2 (scaffolds in the vAM genome were combined using stretches of 100 Ns to denote a gap of unknown length). Since it is possible that PBjelly2 only fills a gap partially, we had to identify the positions of the gaps introduced by AllMaps in the new genome version and checked if they were filled completely or not. If the gap length was reduced, it was extended to have a length of 100 again. This approach produced genome v2.6.

**Figure 1:**
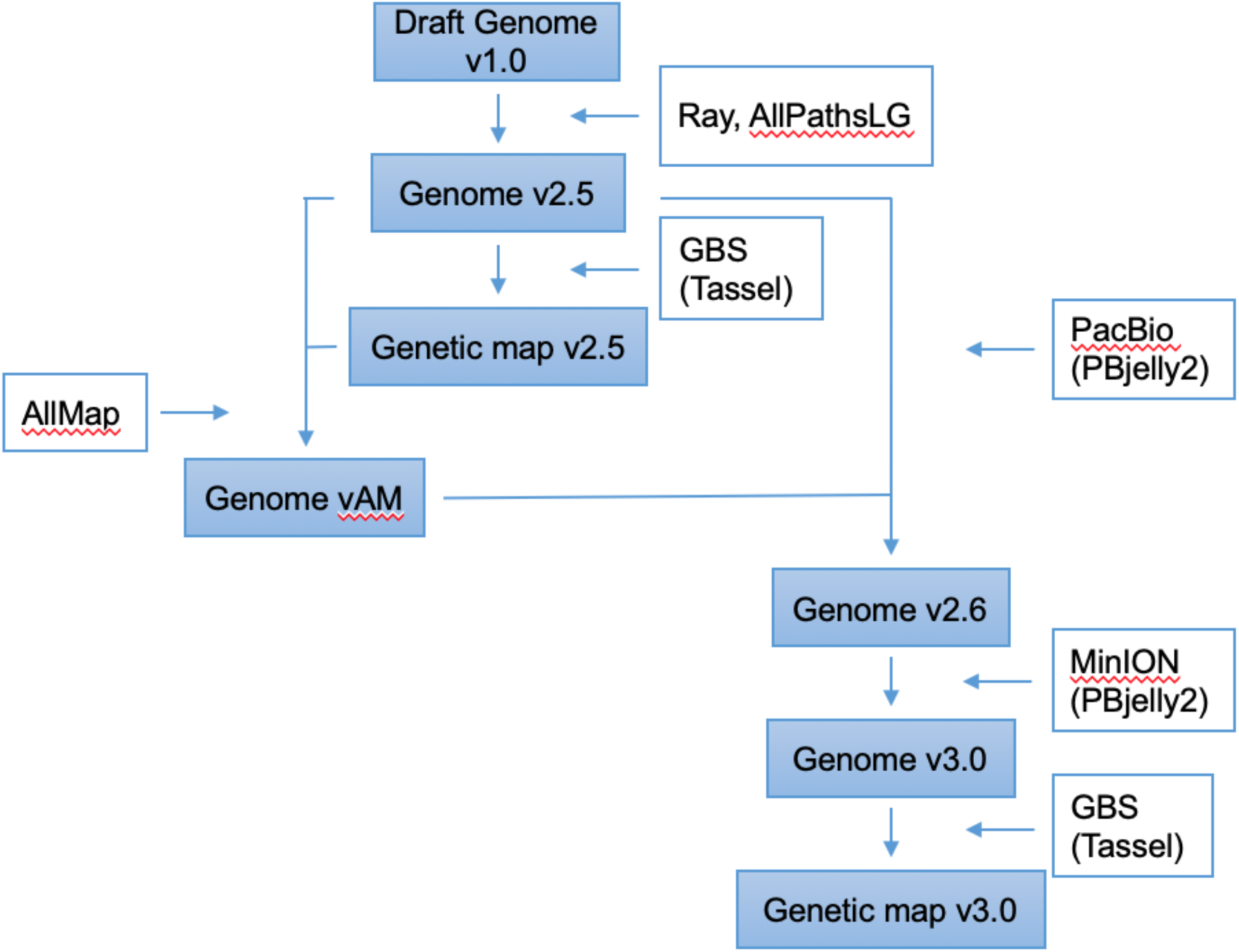
Overview of the analyses performed in this study. In filled boxes are data sets, approaches and companying tools are in open boxes.

**Figure 2:**
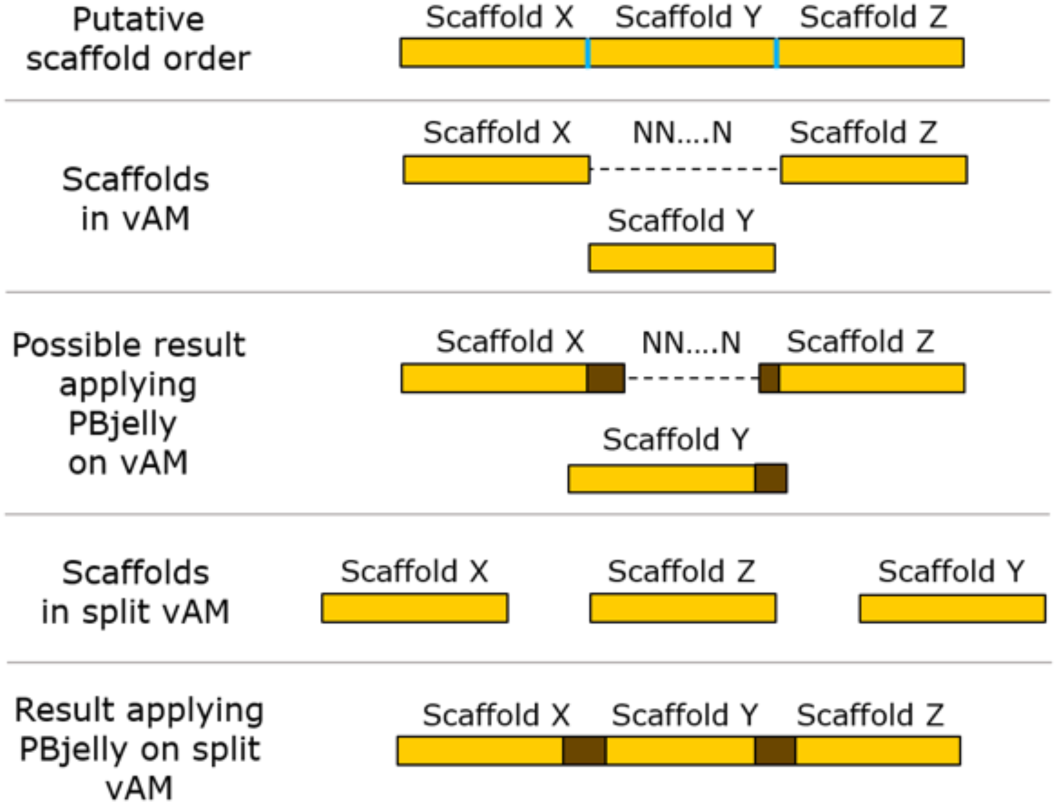
Problem arising from applying PBjelly2 on vAM. Scaffold borders are visualized in blue and extensions of scaffolds introduced by PBjelly2 are shown in brown. Assuming the true order of the scaffolds is shown on top of the figure, but scaffold X and scaffold Z were already combined in the vAM assembly (second bar from top) this could lead to a partial filling of the N-stretch and maybe an extension of scaffold Y. However, PBjelly2 would not be able to place scaffold Y between the two other scaffolds (middle bar). If the scaffolds were thus split again (second bar from bottom), it is possible that the connections are made correctly applying PBjelly2 on the split version (bottom bar). This only visualizes a theoretical case, in this work it appeared every time that scaffold X and Y were connected by PBjelly2 and scaffold Z had to be reconnected afterwards.

#### Genome version 3.0: MinION sequencing incorporation

After improving the genome to v2.6 using the PacBio reads, the same approach was applied for 30,935 MinION TUR reads to obtain the *Ae. arabicum* genome v3.0. The MinION reads were also checked for contamination. The genome version 3.0 is available at https://genomevolution.org/coge/GenomeView.pl?gid=36061.

#### Name convention of *Ae. arabicum* v3.0 genome scaffolds

Scaffolds of genome v3.0 were named and ordered according to their length from long to short. The longest eleven scaffolds were named linkage group (LG) based on the genetic map. Scaffolds which were combined are named csc for concatenated scaffold and the other ones are named sc (scaffold). The v3.0 scaffold names therefore follow the scheme type-number?v2.5 scaffold[.v2.5 scaffold…]. I.e., the scaffold type (LG, CSC, SC), followed by a minus and the number of the scaffold, separated by a blank, followed a list of scaffolds denoting the v2.5 scaffolds or the v3.0 scaffolds. This naming system resulted in a shift in LG order between v2.5 and v3.0 (Supplementary file linkage_group_map.xlsx).

#### Migration of proteins to new genome version

To perform the lift over of the gene models from v2.5 to v3.0, a combination of Gene Model Mapper (GeMoMa) v1.4 [32] and flo (flo - same species annotations lift over pipeline, https://github.com/wurmlab/flo) were used. The results of both programs were concatenated. flo results were preferred over GeMoMa results if the results of the two programs differed, because flo works with alignments on nucleotide level while GeMoMa works with blasting proteins on amino acid level. If a protein could not be lifted completely, it is marked as partial in the resulting GFF (v3.0). A total of 34 genes had to be lifted manually, because they were either not lifted at all or only partially. If an intron could not be lifted, it was added by hand. If an exon or CDS could not be lifted, the new location was deriving from neighboring features which could be migrated to the new genome version. The location was then used to extract the nucleotide sequence from the genome using samtools v1.4 [33]. Only if the sequence was identical to the original sequence extracted from v2.5, the feature was migrated. This check was performed with ClustalW v2.1 [34]. After the migration step, the GFF file was checked and corrected. Genes which did not contain a start or a stop, contained internal stops or whose CDS sequence had a length not dividable by 3 were marked as potential pseudogenes with “pseudo=true”. To check if a gene contains internal stops each of its CDS features was checked individually for having at least one frame which results in no stop codons. Genes which were identical to other genes (start and stop position are equal) or were contained in other genes were removed. If the 3’ CDS of a gene did not contain a stop codon but could be added by extending the CDS by three nucleotides, the CDS was corrected. The lifted genes were classified as shown inTable 2.

**Table 2:** Overview of gene liftover: GFF migration statistics.

|  |  |
| --- | --- |
| Lifted only by flo | 4,346 |
| Lifted only by GeMoMa | 36 |
| Lifted with both programs | 18,177 |
| Manually lifted | 34 |
| Partially lifted | 14 |
| Number of corrected CDS | 10,259 |
| Marked as pseudo | 3,230 |

Most genes could be lifted by flo and GeMoMa. The reason why flo was able to lift more proteins is that GeMoMa works with protein sequences and the program was not able to generate proteins for 20,056 CDS features, either because a gene did not possess a CDS or because of faulty CDS sequences.

#### Name convention of v3.0 genes

Old gene IDs were kept in the note attribute of the genes in the GFF and the linkage group numbers of the genetic map are also noted. The names of the genes were changed into Aa3typeNumberGenenumber: Aa for *Aethionema arabicum*, indicator genome version 3, followed by the type of scaffold, its number and the number of the gene (starting with 1 at the 5’ end),e.g. Aa3LG1G2 or Aa3SC2601G1). For transcript isoforms (splice variants) this locus nomenclature can be extended by the number of the isoform (.X). Version 3.0 of the genome and all gene models are available at https://genomevolution.org/coge/GenomeView.pl?gid=36061.

#### Data Availability Statement

The genome version 2.5 is available at: https://genomevolution.org/coge/GenomeInfo.pl?gid=23428.

The genome version 3.0 is available at: https://genomevolution.org/coge/GenomeView.pl?gid=36061.

For Review: the data (as mapped BAM files) have been made available in CoGe. Account: agr_reviewer Password: GoCoGe’!

https://auth.iplantcollaborative.org/cas4/login?service=https://genomevolution.org/coge/index.pl https://genomevolution.org/coge/GenomeInfo.pl?gid=36061

The GBS unprocessed and processed reads for genome mapping 2.5 and 3.0 are available as supplmental files S1-S4. The linkage group information for both 2.5 and 3.0 genetic maps are available as S5. A summary of the contamination screen is available as S6.

## Results and Discussion

### Reference genome improvement

The published draft version of the *Ae. arabicum* genome utilized the Ray assembler and contained 59,101 scaffolds with an N50 of 115,195bp [9]. Reassembly using the the AllPathsLG assembler and gap-closing using the SOAPdenovo GapCloser tool were used as a starting point for super-scaffolding. This resulted in a reassembly with 3,166 scaffolds, and a scaffold N_50_ of 564,741bp labeled and published as version 2.5 on https://genomevolution.org/coge/. The subsequent genome versions (vAM, v2.6 and v3.0) were obtained using linkage map and long read correction. The quality improvement of the genome is presented as the increase in total number of bases, reduced number of scaffolds and number of gaps, as well as bigger N_50_ and smaller L_50_ parameters (Table 3). In comparison with the starting genome v2.5, the final genome v3.0 has 9% less scaffolds (from 3,166 to 2,883). The overall length of genome v3.0 was extended from 196,005,095 to 203,449,326 bases (17% more) and the number of uncalled based was reduced from 25,768,296 to 13,790,434 (from 13.2% to 6.8%) (Table 3).

**Table 3:** Statistic overview of *Aethionema arabicum* genome versions

| Genome version | Draft | v2.5 | vAM | v2.6 | v3.0 |
| --- | --- | --- | --- | --- | --- |
| # Bases |  | 196,005,095 | 196,022,695 | 203,150,143 | 203,449,326 |
| # Scaffolds | 59,101 | 3,166 | 2,990 | 2,895 | 2,883 |
| # Scaffolds containing Ns |  | 1,910 | 1,734 | 1,542 | 1,539 |
| # Ns |  | 25,768,296 | 25,785,896 | 13,946,922 | 13,790,434 |
| N50 | 115,195 | 564,741 | 10,141,718 | 10,328,388 | 10,328,388 |
| L50 |  | 56 | 9 | 9 | 9 |

### Genome improvement using long reads

#### Read mapping efficiency

The results of the mapping of the reads to the genome using PBjelly2 are summarized in Table 4. Almost the same number of PBjelly2 reads were mapped to genome v2.5 and vAM. However, it was important to apply PBjelly2 on both genomes in order to find scaffold connections which were not possible due to a combination of certain scaffolds in vAM (see supplementary file combination_comparison_pbjelly_for_v2.5_vs_pbjelly_for_vGM.ods for details). The genome v2.6 resulted by improving the split vAM genome using PBjelly2, reconnecting scaffolds and resizing gaps if needed. We also compared the results for improving v2.5 with vAM, but there were no new scaffold connections which were missed by improving the v2.6 version, so we did not perform a split step for improving the genome using the MinION reads. The mapping efficiency for the MinION reads is lower than for the PacBio reads due to a contamination of the reads (see Methods for details). There are 5.9% more MinION reads which were mapped to v2.6 than v2.5, demonstrating that the changes done to the genome are supported by the very long reads.

**Table 4:** Mapping efficiency of PBjelly2’s mapping step. The percentages in brackets give the percentage of the total number of reads (CYP, TUR or CYP + TUR). The line “# covered scaffolds” gives the number of scaffolds in which at least one read was mapped. Here, the number in brackets gives the percentage of the total number of scaffolds.

| **Setup** | **PacBio vs. v2.5** | **PacBio vs. vAM** | **MinION vs. v2.5** | **MinION vs. v2.6** |
| --- | --- | --- | --- | --- |
| **# mapped TUR reads** | 198,675<br>(86.9%) | 198,629<br>(86.9%) | 14,098 (45.6%) | 15,886 (51.4%) |
| **# mapped CYP reads** | 131,976<br>(86.6%) | 131,942<br>(86.6%) | - | - |
| **# total reads mapped** | 330,651<br>(86.8%) | 330,571<br>(86.8%) | 14,098 (45.6%) | 15,886 (51.4%) |
| **# unmapped reads** | 50,371 (13.2%) | 50,451 (13.2%) | 16,837 (54.4%) | 15,049 (48.6%) |
| **# scaffolds input genome** | 3,166 | 2,990 | 3,166 | 2,895 |
| **# covered scaffolds** | 2,971 (93.8%) | 2,804 (93.8%) | 1,689 (53.3%) | 1,429 (49.4%) |

#### The effect of PBjelly2 runs applied to the different assembly versions

Most scaffolds were combined in the vAM which resulted in the L_50_ valuelowered from 56 to 9 and the N_50_ value almost doubled. PBjelly2 was not as good as using the genetic map in combining scaffolds. The increase of the N_50_ value in case of the PBjelly2 result (using the PacBio reads for improving scaffolds) results from more improvements of the shorter scaffolds. Comparing the PBjelly2 result for applying the PacBio reads to v2.5 and vAM shows that the reduction of scaffold number and increase of number of bases in the genome is similar (Table 5). MinION reads could also be used for v2.5 assembly improvement, however the results were not as good as for using PacBio reads, due to a much smaller number of reads. Improving the genome v2.6 with the MinION reads is also possible, but the improvement is not as good as for v2.5. This demonstrates that connections done using the PacBio reads are also supported by MinION reads.

**Table 5:** Overview of the PBjelly2 result statistics for the different setups.

| Setup | v2.5 | PacBio vs. v2.5 | MinION vs. v2.5 |
| --- | --- | --- | --- |
| # scaffolds | 3,166 | 3,066 | 3,123 |
| # bases | 196,005,095 | 203,024,676 | 196,600,700 |
| N50 | 564,741 | 542,490 | 564,741 |
| L50 | 56 | 58 | 56 |
| Setup | vAM | PacBio vs. vAM |  |
| # scaffolds | 2,990 | 2,905 |  |
| # bases | 196,022,695 | 203,137,854 |  |
| N50 | 10,141,718 | 10,314,234 |  |
| L50 | 9 | 9 |  |
| Setup | v2.6 |  | MinION vs. v2.6 |
| # scaffolds | 2,895 |  | 2,886 |
| # bases | 203,150,143 |  | 203,450,934 |
| N50 | 10,328,388 |  | 10,323,234 |
| L50 | 9 |  | 9 |

Comparison of values for the different genome versions with the values for the PBjelly2 output is shown. The PBjelly2 outputs are denoted in the form “read type” vs. “genome version” to show which reads were used to improve which version of the genome. The result for PacBio vs. vAM was the basis for v2.6 and MinION vs. v2.6 was the basis for v3.0.

While PBjelly2 does not do as good a job as the genetic map approach in connecting scaffolds, its power is revealed by the gap filling. In genome v2.5 a total of 1,910 scaffolds contained uncalled bases. This number was reduced to 1,711 scaffolds (by 7.3%; Table 6) using PBjelly2 with PacBio reads. The exact number of uncalled bases in the v2.5 *A. arabicum* genome was 25,768,296 (Table 6). In the PBjelly2 result only 13,940,203 Ns (Table 6) are left, a reduction by 45.9%. Comparing this with the PBjelly2 result for the improvement of the vAM genome using PacBio reads (Table 6), more gaps were removed from the connected genome. The number of scaffolds containing Ns was reduced by 10.8% and the overall number of Ns was reduced by 50.0%, while the overall percentage of Ns in the genome remained the same in the two results. Due to the small number of MinION reads, the improvement of the assembly is less pronounced.

**Table 6:**
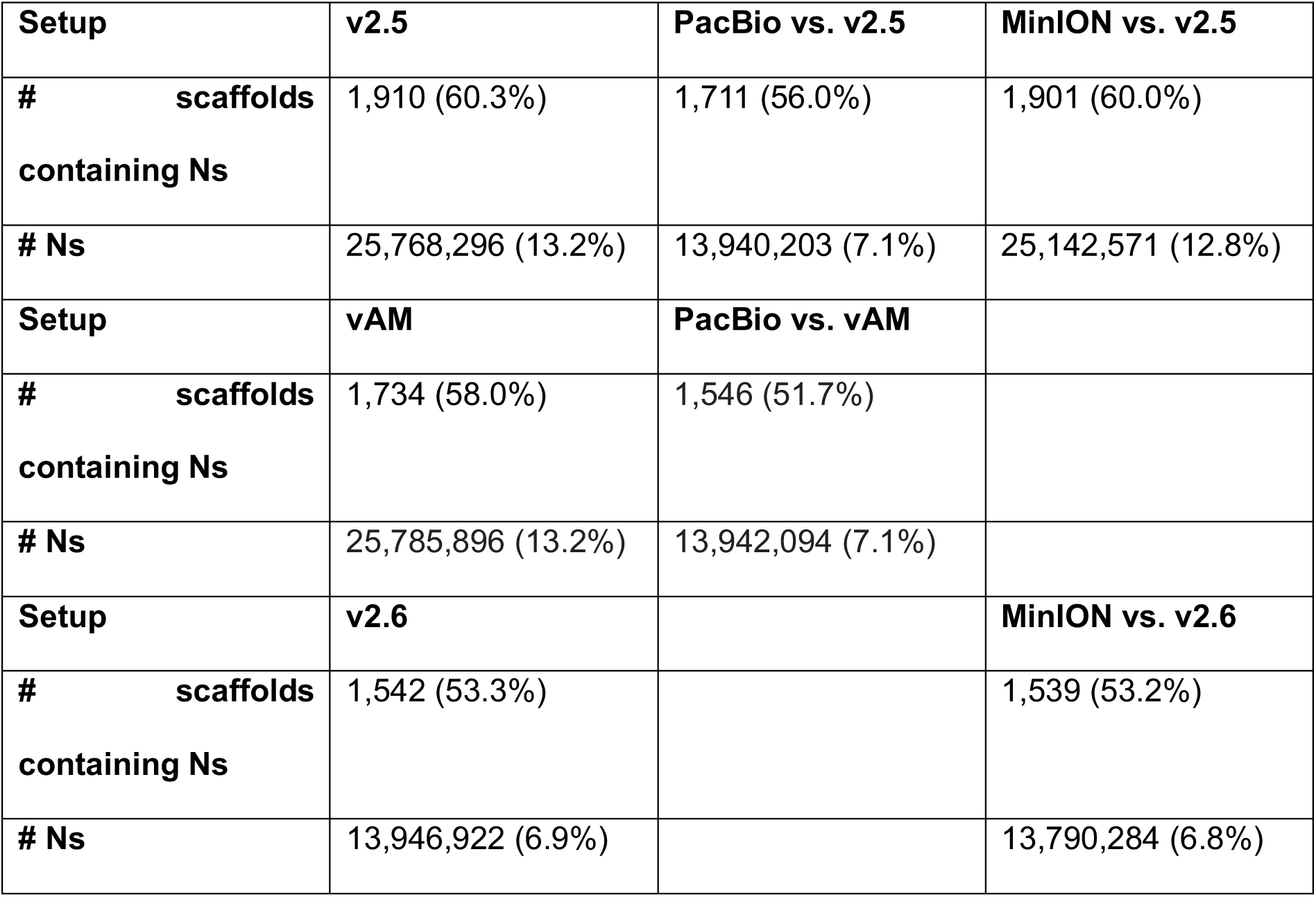
Gap/N analysis of different genome versions.

| Setup | v2.5 | PacBio vs. v2.5 | MinION vs. v2.5 |
| --- | --- | --- | --- |
| # scaffolds containing Ns | 1,910 (60.3%) | 1,711 (56.0%) | 1,901 (60.0%) |
| # Ns | 25,768,296 (13.2%) | 13,940,203 (7.1%) | 25,142,571 (12.8%) |
| Setup | vAM | PacBio vs. vAM |  |
| # scaffolds containing Ns | 1,734 (58.0%) | 1,546 (51.7%) |  |
| # Ns | 25,785,896 (13.2%) | 13,942,094 (7.1%) |  |
| Setup | v2.6 |  | MinION vs. v2.6 |
| # scaffolds containing Ns | 1,542 (53.3%) |  | 1,539 (53.2%) |
| # Ns | 13,946,922 (6.9%) |  | 13,790,284 (6.8%) |

This table gives an overview of the number of Ns in the different genome versions and the PBjelly2 results.. For the number of scaffolds containing Ns, the percentage is given relative to the total number of scaffolds is given in brackets. For the number of Ns, the percentage is relative to the total number of bases in the respective assembly.

### Migration of proteins to new genome version

The genome v2.5 harbors 23,594 annotated protein coding genes. Eight of them could not be lifted because they were located next to a gap in the genome. Since it is possible that PBjelly2 changes the sequences around gaps, the sequences of the genes were not identical anymore and the programs were therefore not able to migrate some genes from one assembly version to another. We checked the expression of the genes which could not be lifted using Illumina RNA-seq data representing several developmental stages (data not shown) and found that all of them had almost no expression, as a result they were not lifted manually. In addition to some unlifted genes, there were 17 genes which could be lifted only partially due to the same reason. All the other 23,569 genes could be lifted. A set of 579 genes were removed due to being identical with other genes, and 140 genes were removed because they were located completely in another gene. A total of 1,202 genes have no starting methionine, 2,055 have no stop, 132 genes contain internal stops and for 1,019 genes the length of the CDSs is not dividable by three. In the end 19,363 genes were lifted which were not marked as potential pseudogene or partial.

We find that that start point (assembly version) for improvement is not relevant. PBjelly2 is able to make more improvements using the PacBio data than with the MinION data due to the higher number of PacBio reads. The number of added bases per read is higher for PacBio than MinION reads (18.71 vs. 9.67) and also the number of removed Ns is higher (31.07 vs. 5.06) while the MinION reads lead to more scaffold connections per read (2.49 × 10^−4^ vs. 3.88 × 10^−4^ connected scaffolds). Using the MinION reads for improvement makes only a few changes, but they show that they support the changes which were made to the genome using the PacBio data. Since the *Ae. arabicum* genome was almost not contaminated at all, only three small scaffolds had to be removed. Gene models were filtered for multiple genes and genes contained in other genes. If a problem with a gene was found, it was marked in the resulting GFF file. We note that there are gene models which are probably not correct and need to be fixed in future studies.

The combination of genetic mapping and long reads significantly improved the structure of the assembly, reducing the total scaffold number and decreasing the number of gaps.

### Genetic map of *Aethionema arabicum*

#### Genetic map v2.5

##### SNP calling

A GBS approach [35] was used to generate genetic variation data for genetic mapping. Illumina sequencing of the parental lines and the RILs resulted in 442,101,405 raw reads after quality filtering. Using the TASSEL package [19] to match sequence tags to markers, 160,379 SNPs could be called based on genome v2.5. SNPs identified through the GBS method often take the form of many SNP ‘islands’, where a multitude of SNPs is present over only a few kbp of sequence with the same states across individuals. This makes genetic mapping difficult as it results in a very large number of markers that are mostly redundant. We reduced this SNP amount using a sliding window approach collapsing a group of SNPs that all have the same states across individuals into one single marker over windows of 10 kbp, thus the bigger the scaffold the more selected SNPs. This, together with filtering non-informative SNPs (missing data in more than 30% many individuals) resulted in a core group of markers of 5,428 SNPs.

##### Genetic map calculation

We used JoinMap 4.1 to calculate the genetic linkage map for the *Ae. arabicum* RIL population. For map v2.5, we first checked the marker similarity among the initial set of 5,428 SNPs that were obtained from GBS based on genome v2.5 by a pairwise comparison. SNPs that were highly similar (higher than 90%) were represented by one marker, which refined the number of markers to 1,818. Grouping was selected at a LOD threshold of 9.0, which led to the grouping of the expected 11 Linkage Groups (LGs) (Figure 3). We further optimized each LG to avoid inflation of the map distance due to saturating SNPs using a Maximum likelihood model. Markers that were not more than 0.1 cM away were also eliminated. As a result, a final set of 746 SNP markers was used on 11LGs. Out of 11 LGs, there are three LGs (4, 7 and 11) containing cluster of segregation distorted SNPs (more than 50% of SNPs, supplemental file linkage_group_map.xlsx).

**Figure 3:**
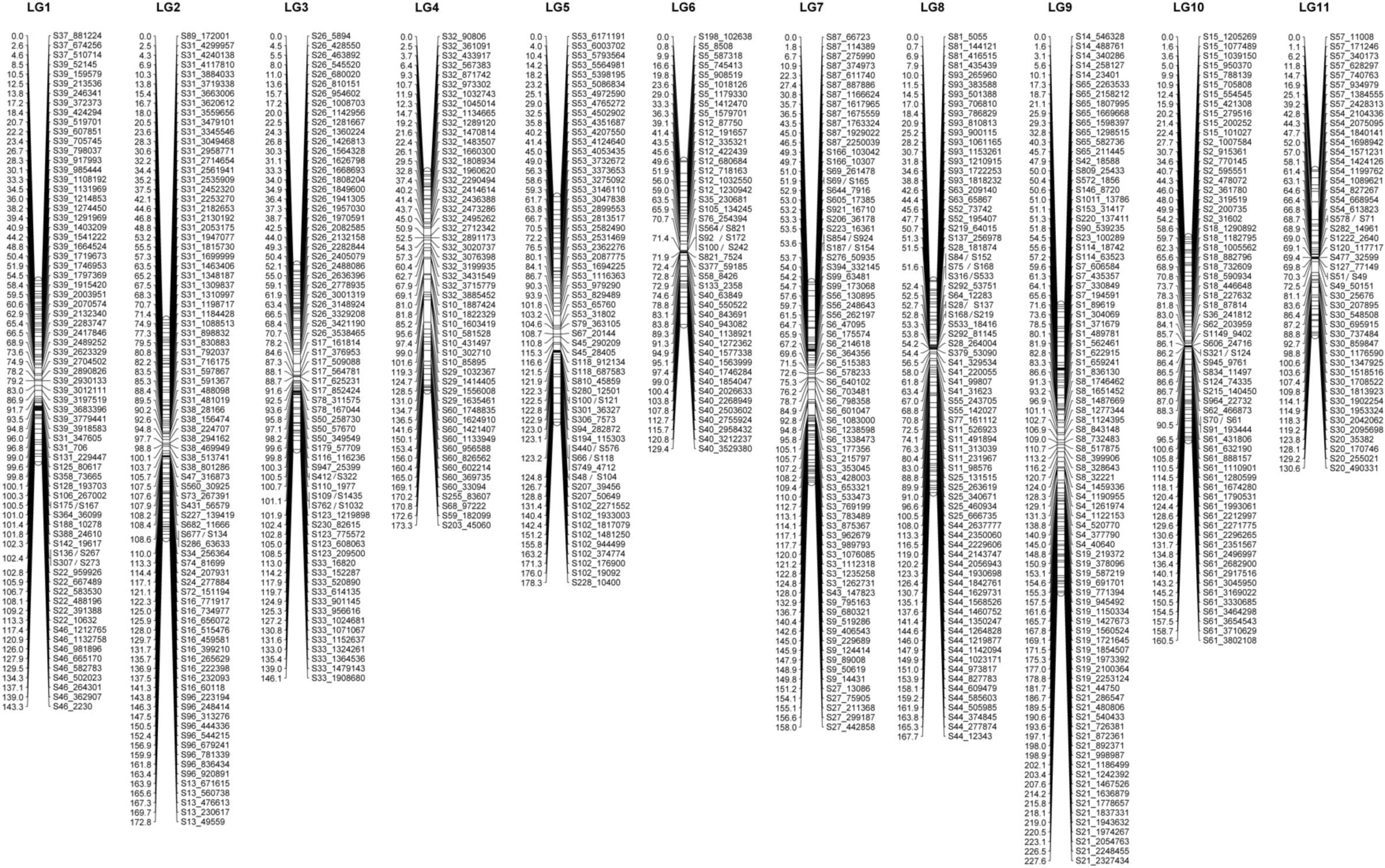
*Aethionema arabicum* genetic map v2.5. Genetic map version 2.5 consists of eleven linkage groups. On each linkage group, genetic distance in cM is present on the left and SNP markers on the right.

The *Ae. arabicum* genetic map v2.5 consists of 11 LGs with average distance size of 162.5 cM, covered by 746 SNP markers with average of 67 markers per LG. The average marker spacing was 2.4 cM, equivalent to approximately 169 kbp. The centromere is suggested by the high density of SNP markers within a small genetic distance (e.g. a low recombination frequency). These markers typically belong to relatively small scaffolds, consistent with a high-repetitive DNA content, where only a few SNPs were called. LG4 centromere is located at the end of the linkage, suggesting an assembly problem or that LG4 is a telocentric chromosome.

Overall the markers are distributed relatively dense and even in v2.5, with the biggest gap smaller than 18 cM. SNPs that reside in the same scaffold were in agreement among each other on the direction of their scaffold.

#### Genetic map v3.0

The procedure to build genetic map v3.0 was similar to v2.5. SNP calling was performed based on genome v3.0 instead of v2.5 resulting in a raw set of 141,914 SNPs. After similar quality control strategy as for v2.5, we construct v3.0 with a core set of 632 SNPs (Figure 4). The 11 LGs were maintained with the total size of~ 1945 cM, average marker distance of 3.1 cM. This inflation of genetic distance in genetic map v3.0 compared with v2.5 can be explain by the newly retrieved SNPs due to resolved Ns in the genome. These new SNPs are mainly located in the centromeric regions. In general, SNP order and orientation in LGs are in agreement with map v2.5 (Figure 5). We have made adjusted linkage group order in map v3.0 compared with map v2.5 (linkage_group_map.xlsx). Based on the size of the group, the biggest one is LG 1, and the smallest is LG 11 (Figure 4).

**Figure 4:**
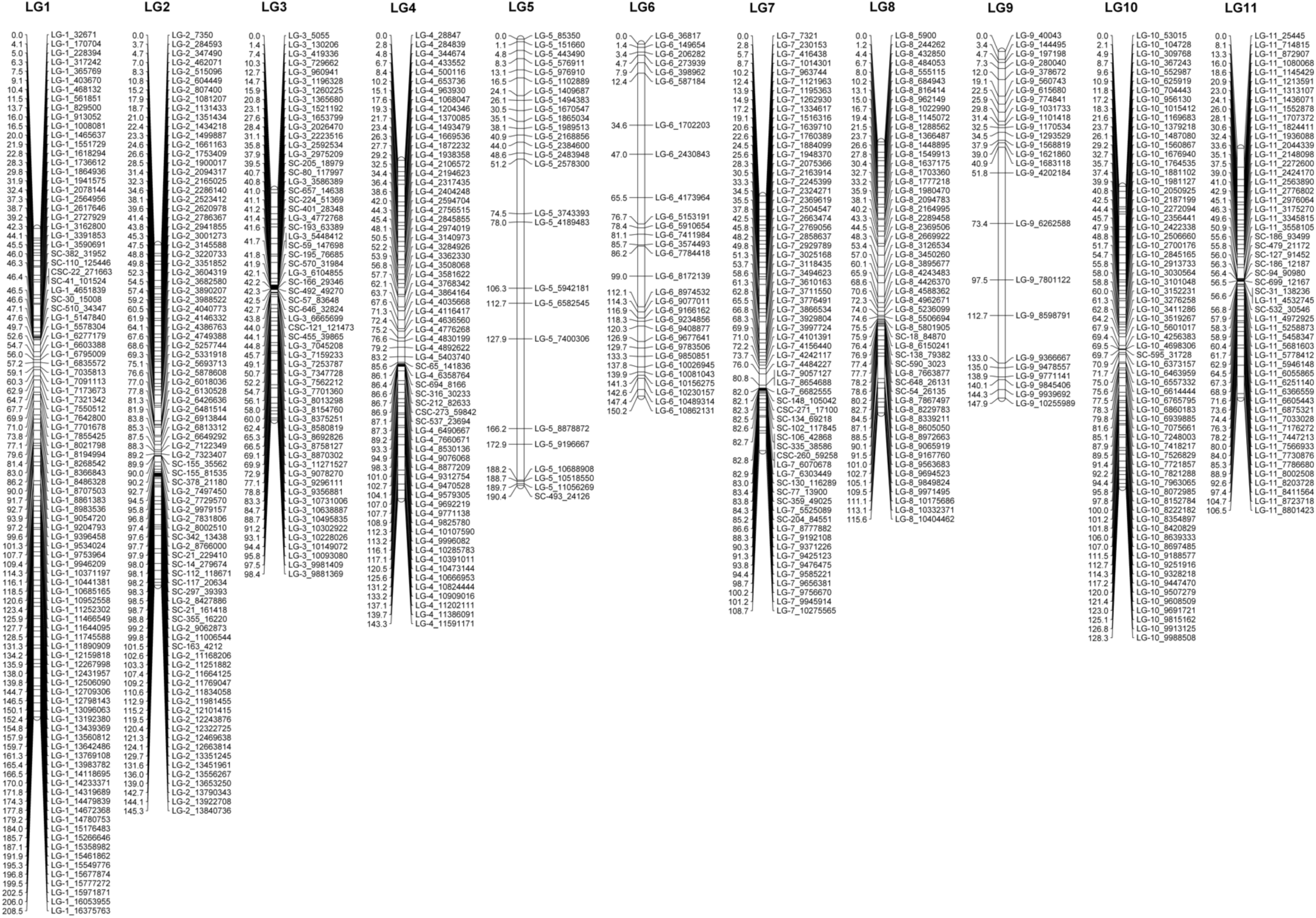
*Aethionema arabicum* genetic map v3.0. Genetic map version 3.0 consists of eleven linkage groups. On each linkage group, genetic distance in cM is present on the left and SNP markers on the right.

**Figure 5:**
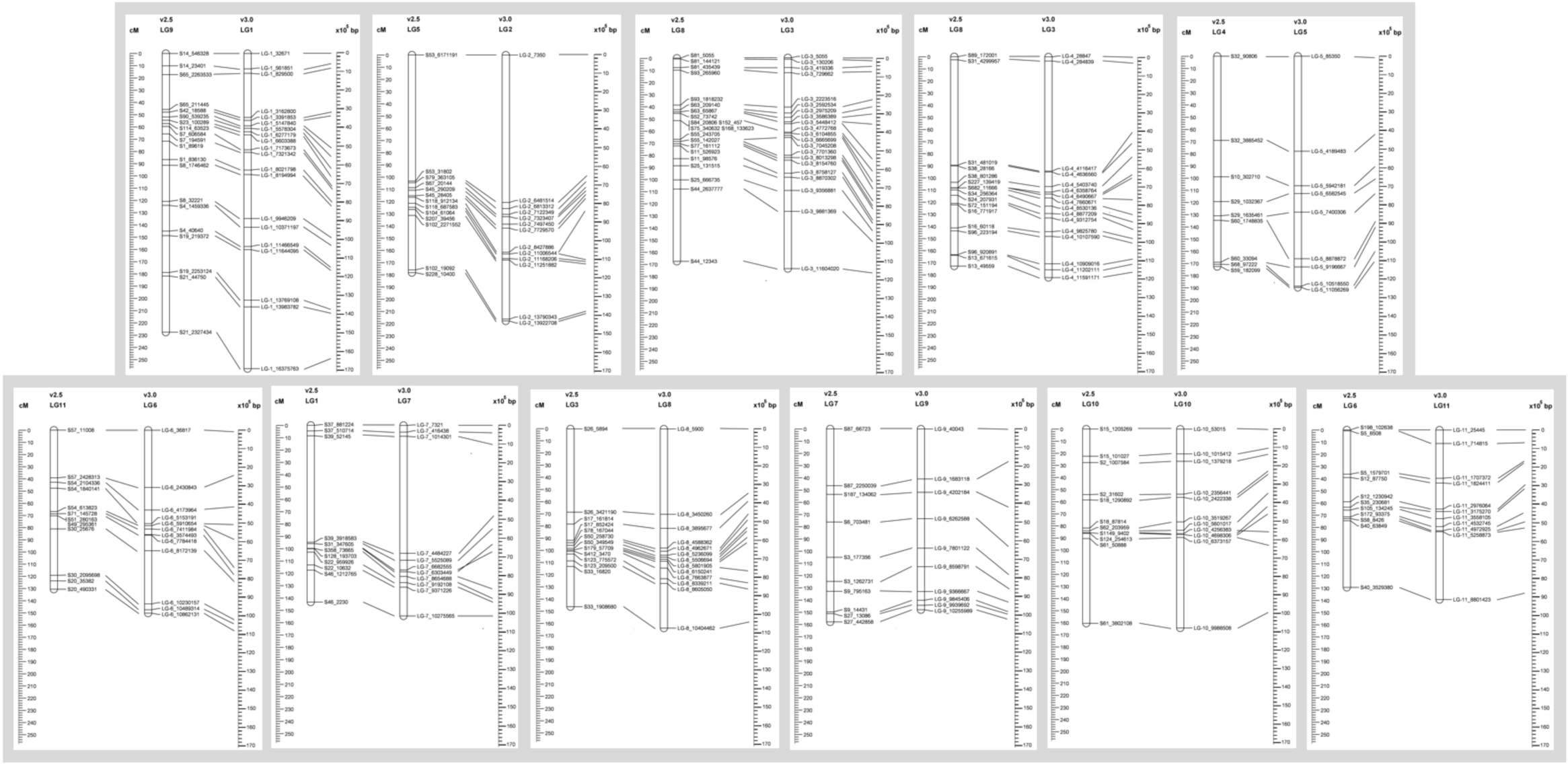
The alignment of genetic map v2.5, v3.0 and physical map. The alignment of the genetic map v2.5 and v3.0 were based on relative SNPs. The left ruler indicates genetic distance in cM and the right indicates physical distance in bp according to genome v3.0

However, there is a significant difference between v3.0 and v2.5 at three LGs that harbor clusters of segregation distorted SNPs, LG 5, 6, and 9 (equivalent LG 4, 11 and 7 in v2.5, respectively): the reduced number of markers as well as the increased marker distance (Figure 3 and 4). In order to maintain 11 LGs and certain degree of newly called SNP incorporation, we had to reduced number of distorted markers in those LGs in v3.0, as a result the dearth of markers was observed (see supplemental file. linkage_group_map.xlsx)

## Conclusions and potential implications

*Aethionema* is becoming an outgroup model for the Brassicaceae core. Studies on its genome, relevant life-history traits and their evolution rely on genome and genetic map resources. Thus, this work helps pave the way for future research of the Brassicaceae family. We have provided an advanced version of the *Aethionema arabicum* genome and its first genetic map, which allows for pseudochromosome construction needed for analysis of genome evolution. Finally, quantitative trait loci (QTL) mapping for the wide range of traits in *Aethionema* (e.g. flowering time, plant fitness, chemical defense and heteromorphism) will be greatly enabled by this genetic map.

## Declarations

None of the authors have any competing interests in the manuscript.

## List of abbreviations

BAC: Bacterial Artificial Chromosome
BS-seq: Bisulfite sequencing
ChIP-seq: Chromatin ImmunoPrecipitation DNA-Sequencing
CYP: Cyprus ecotype
FPC: Finger Printed Contigs
GFF: Generic/General Feature Format
LG: Linkage group
MinION: Oxford Nanopore MinION
PacBio: Pacific Biosciences
QTL: Quantitative trait loci
RIL: Recombinant Inbred Line
SNP: Single Nucleotide Polymorphism
TUR: Turkey ecotype
WGP: Whole Genome Profiling

## Authors’ contributions

TPN and ES designed the GBS experiment and performed the genetic map analyses. SM contributed to the development of the RIL population. AP did the initial genome assemblies. CM analysed the PacBio data, used the long-read data for super-scaffolding and carried out the liftover of gene models to v3.0. FBH analysed the MinION data and carried out the contamination check. SAR supervised the work done by CM and FBH and conceived of this part of the work. EvdB performed GBS SNP calling. TPN, CM, SAR and ES wrote the manuscript with contributions by all authors.

## Acknowledgements

We thank Dr. Elio Schijlen for technical support on Genotyping by Sequencing technology, Dr. Lidija Berke for her help on running AllMaps and Marco Göttig, Rabea Meyberg, Christopher Grosche and Manuel Hiss for their help with long read sequencing. We also thank the other members of the SeedAdapt Consortium and Dr. Laurie Grandont for fruitful discussions about the work.

## References

1. Warwick SI and Al-Shehbaz IA. Brassicaceae: Chromosome number index and database on CD-Rom. Plant Syst Evol. 2006;259 2-4:237–48. doi:10.1007/s00606-006-0421-1.

2. Al-Shehbaz IA, Beilstein MA and Kellogg EA. Systematics and phylogeny of the Brassicaceae (Cruciferae): an overview. Plant Syst Evol. 2006;259 2-4:89–120. doi:10.1007/s00606-006-0415-z.

3. Beilstein MA, Al-Shehbaz IA, Mathews S and Kellogg EA. Brassicaceae phylogeny inferred from phytochrome A and ndhF sequence data: tribes and trichomes revisited. Am J Bot. 2008;95 10:1307–27. doi:10.3732/ajb.0800065.

4. Franzke A, Lysak MA, Al-Shehbaz IA, Koch MA and Mummenhoff K. Cabbage family affairs: the evolutionary history of Brassicaceae. Trends Plant Sci. 2011;16 2:108–16. doi:10.1016/j.tplants.2010.11.005.

5. Imbert E. Ecological consequences and ontogeny of seed heteromorphism. Perspect Plant Ecol. 2002;5 1:13–36. doi:Doi 10.1078/1433-8319-00021.

6. Lenser T, Graeber K, Cevik ZS, Adiguzel N, Donmez AA, Grosche C, et al. Developmental control and plasticity of fruit and seed dimorphism in Aethionema arabicum. Plant Physiol. 2016;172 3:1691–707. doi:10.1104/pp.16.00838.

7. Mohammadin S, Peterse K, van de Kerke SJ, Chatrou LW, Donmez AA, Mummenhoff K, et al. Anatolian origins and diversification of Aethionema, the sister lineage of the core Brassicaceae. Am J Bot. 2017; doi:10.3732/ajb.1700091.

8. Bibalani GH. Investigation on flowering phenology of Brassicaceae in the Shanjan region Shabestar district, NW Iran (usage for honeybees).. Annals of Biological Research. 2012; 6: 1958 –68.

9. Haudry A, Platts AE, Vello E, Hoen DR, Leclercq M, Williamson RJ, et al. An atlas of over 90,000 conserved noncoding sequences provides insight into crucifer regulatory regions. Nature genetics. 2013;45 8:891–U228. doi:10.1038/ng.2684.

10. Boisvert S, Laviolette F and Corbeil J. Ray: simultaneous assembly of reads from a mix of highthroughput sequencing technologies. Journal of computational biology : a journal of computational molecular cell biology. 2010;17 11:1519–33. doi:10.1089/cmb.2009.0238.

11. Gnerre S, MacCallum I, Przybylski D, Ribeiro FJ, Burton JN, Walker BJ, et al. High-quality draft assemblies of mammalian genomes from massively parallel sequence data. P Natl Acad Sci USA. 2011;108 4:1513–8. doi:10.1073/pnas.1017351108.

12. Tang H, Zhang X, Miao C, Zhang J, Ming R, Schnable JC, et al. ALLMAPS: robust scaffold ordering based on multiple maps. Genome Biol. 2015;16:3. doi:10.1186/s13059-014-0573-1.

13. English AC, Richards S, Han Y, Wang M, Vee V, Qu J, et al. Mind the gap: upgrading genomes with Pacific Biosciences RS long-read sequencing technology. Plos One. 2012;7 11:e47768. doi:10.1371/journal.pone.0047768.

14. Luo RB, Liu BH, Xie YL, Li ZY, Huang WH, Yuan JY, et al. SOAPdenovo2: an empirically improved memory-efficient short-read de novo assembler. Gigascience. 2012;1 1:18. doi:10.1186/2047-217x-1-18.

15. Kent WJ, Sugnet CW, Furey TS, Roskin KM, Pringle TH, Zahler AM, et al. The human genome browser at UCSC. Genome Res. 2002;12 6:996–1006. doi:10.1101/gr.229102.

16. Mohammadin S, Wang W, Liu T, Moazzeni H, Ertugrul K, Uysal T, et al. Genome-wide nucleotide diversity and associations with geography, ploidy level and glucosinolate profiles in Aethionema arabicum (Brassicaceae). Plant Syst Evol. 2018;304 5:619–30. doi:10.1007/s00606-018-1494-3.

17. Doyle J. DNA Protocols for Plants. In: Hewitt GM, Johnston AWB and Young JPW, editors. Molecular Techniques in Taxonomy. Springer-Verlag Berlin Heidelberg: Springer Berlin Heidelberg; 1991. p. 283–93.

18. Elshire RJ, Glaubitz JC, Sun Q, Poland JA, Kawamoto K, Buckler ES, et al. A Robust, Simple Genotyping-by-Sequencing (GBS) Approach for High Diversity Species. Plos One. 2011;6 5:e19379. doi:10.1371/journal.pone.0019379.

19. Glaubitz JC, Casstevens TM, Lu F, Harriman J, Elshire RJ, Sun Q, et al. TASSEL-GBS: A High Capacity Genotyping by Sequencing Analysis Pipeline. Plos One. 2014;9 2:e90346. doi:10.1371/journal.pone.0090346.

20. Li H and Durbin R. Fast and accurate long-read alignment with Burrows-Wheeler transform. Bioinformatics. 2010;26 5:589–95. doi:10.1093/bioinformatics/btp698.

21. Stam P. Construction of Integrated Genetic-Linkage Maps by Means of a New Computer Package - Joinmap. Plant J. 1993;3 5:739–44. doi:DOI 10.1111/j.1365-313X.1993.00739.x.

22. Ooijen JWV. JoinMap ® 4, Software for the calculation of genetic linkage maps in experimental populations. Wageningen, Netherlands: Kyazma B.V., 2006.

23. Coordinators NR. Database Resources of the National Center for Biotechnology Information. Nucleic acids research. 2017;45 D1:D12–D7. doi:10.1093/nar/gkw1071.

24. Huson DH, Beier S, Flade I, Gorska A, El-Hadidi M, Mitra S, et al. MEGAN community edition - Interactive exploration and analysis of large-scale microbiome sequencing data. PLoS computational biology. 2016;12 6:e1004957. doi:10.1371/journal.pcbi.1004957.

25. Hiss M, Meyberg R, Westermann J, Haas FB, Schneider L, Schallenberg-Rudinger M, et al. Sexual reproduction, sporophyte development and molecular variation in the model moss Physcomitrella patens: introducing the ecotype Reute. Plant J. 2017;90 3:606–20. doi:10.1111/tpj.13501.

26. Dellaporta SL, Wood J and Hicks JB. A plant DNA minipreparation: Version II. Plant Molecular Biology Reporter 14:19–21.

27. Watson M, Thomson M, Risse J, Talbot R, Santoyo-Lopez J, Gharbi K, et al. poRe: an R package for the visualization and analysis of nanopore sequencing data. Bioinformatics. 2015;31 1:114–5. doi:10.1093/bioinformatics/btu590.

28. Wu TD and Nacu S. Fast and SNP-tolerant detection of complex variants and splicing in short reads. Bioinformatics. 2010;26 7:873–81. doi:10.1093/bioinformatics/btq057.

29. Lang D, Ullrich KK, Murat F, Fuchs J, Jenkins J, Haas FB, et al. The Physcomitrella patens chromosome-scale assembly reveals moss genome structure and evolution. Plant J. 2018;93 3:515–33. doi:10.1111/tpj.13801.

30. Chaisson MJ and Tesler G. Mapping single molecule sequencing reads using basic local alignment with successive refinement (BLASR): application and theory. BMC bioinformatics. 2012;13:238. doi:10.1186/1471-2105-13-238.

31. Bolger AM, Lohse M and Usadel B. Trimmomatic: a flexible trimmer for Illumina sequence data. Bioinformatics. 2014;30 15:2114–20. doi:10.1093/bioinformatics/btu170.

32. Keilwagen J, Wenk M, Erickson JL, Schattat MH, Grau J and Hartung F. Using intron position conservation for homology-based gene prediction. Nucleic acids research. 2016;44 9:e89. doi:10.1093/nar/gkw092.

33. Li H, Handsaker B, Wysoker A, Fennell T, Ruan J, Homer N, et al. The Sequence Alignment/Map format and SAMtools. Bioinformatics. 2009;25 16:2078–9. doi:10.1093/bioinformatics/btp352.

34. Larkin MA, Blackshields G, Brown NP, Chenna R, McGettigan PA, McWilliam H, et al. Clustal W and Clustal X version 2.0. Bioinformatics. 2007;23 21:2947–8. doi:10.1093/bioinformatics/btm404.

35. Rowan BA, Patel V, Weigel D and Schneeberger K. Rapid and inexpensive whole-genome genotyping-by-sequencing for crossover localization and fine-scale genetic mapping. G3. 2015;5 3:385–98. doi:10.1534/g3.114.016501.

